# Effects of ERK1/2 Signaling on Cell Cycle Regulation by the Tuberin-Cyclin B1 Complex

**DOI:** 10.64898/2026.06.03.729845

**Authors:** Adam Pillon, Ali Nadi, Jeffery Martin, Miranda A. Hanna, Elizabeth Fidalgo da Silva, Lisa A. Porter

## Abstract

How cells balance growth (cell size) and division (cell number) requires a complex interplay between response to external signals including growth factors, nutrient availability and metabolic cues, along with regulation of the cell cycle. The protein Tuberin (gene *TSC2*) is a critical regulator of these decisions. In a complex with the protein Hamartin, Tuberin functions as a negative regulator of the Target of Rapamycin (mTOR) pathway, preventing excessive growth under unfavorable conditions. However, how this growth pathway connects to decisions to progress through the G2 phase of the cell cycle and permit cell division is still unclear. In this study, we show that post-translational modification of Tuberin by the **<u>E</u>**xtracellular Signal-**<u>R</u>**egulated **<u>K</u>**inase (ERK) pathway abrogates binding between Tuberin and the mitotic cyclin, Cyclin B1. This causes an increase in mitotic cells, due to an unregulated G2/M transition, increasing the proliferation rate. Our work shows a novel role of Tuberin in cell cycle regulation by growth and mitogenic factors independent of mTOR regulation.

## Introduction

The **<u>E</u>**xtracellular Signal-**<u>R</u>**egulated **<u>K</u>**inase 1/2 (ERK1/2) functions as one key component of the **<u>M</u>**itogenic-**<u>A</u>**ctivated **<u>P</u>**rotein **<u>K</u>**inase (MAPK) pathway to phosphorylate proteins responsible for the maintenance of homeostasis and response to change required during development and/or times of stress or changes in the external environment. ERK activation mediates signals that promote cell growth and proliferation, differentiation, survival and apoptosis. Dysregulation of the ERK1/2 pathway has been observed in many diseases, including cancer and neurological disorders [1].

The tumour suppressor protein Tuberin (gene *TSC2*) is one of the main players in the regulation of cell growth and proliferation. This protein forms the **<u>T</u>**uberous **<u>S</u>**clerosis **<u>C</u>**omplex (TSC) together with the proteins Hamartin (gene *TSC1*) and TBC1D7 (gene *TBC1D7*) [2]. About two thirds of the pathogenic mutations that occur in TSC are found in the *TSC2* gene with only about 40% of the mutations occurring in *TSC1* [3,4]. Patients harbouring these mutations develop hamartomas in a plethora of organs, including the brain, lung, heart, kidney and skin. The phenotypic variance of the disorder means that some patients experience small skin fibromas, and some patients can exhibit subependymal giant cell astrocytoma (SEGA) that are implicated in neuropathologies including seizures, autism spectrum disorders, and other forms of neurological development disorders [5].

Both Tuberin and Hamartin are post-translationally modified by phosphorylation from a host of critical signaling pathways in the cell. The canonical function of the TSC is highly studied and is known to be mediated through the **<u>G</u>**TPase **<u>A</u>**ctivating **<u>P</u>**rotein (GAP) domain on the C-terminal of Tuberin. This GAP domain controls the activity of the small G-protein, **<u>R</u>**as-**<u>H</u>**omolog **<u>E</u>**nriched in **<u>B</u>**rain (Rheb) and subsequently acts as an “on-off switch” for the **<u>m</u>**ammalian **<u>T</u>**arget of **<u>R</u>**apamycin (mTOR) or protein synthesis pathway [6]. Tuberin is phosphorylated by a host of kinases including Akt and ERK1/2 which results in an inactivation of Tuberin’s GAP function, thereby permitting increased protein synthesis [7–10]. It is also phosphorylated by AMPK in low nutrient conditions leading to inhibition of mTOR [11,12].

Tuberin and Hamartin directly interact and regulate important cell cycle regulatory proteins. Hamartin interacts with Plk1 and is phosphorylated by the G2 CDK, CDK1 [13,14]. Tuberin can regulate the G1/S CDK, CDK2, by increasing the nuclear accumulation of the CDK2 inhibitor p27 [15]. Tuberin can also directly interact with the G2 cyclins, Cyclin A and Cyclin B1, independent of mTOR activity [16,17]. Tuberin-Cyclin B1 binding lengthens G2 phase of the cell cycle and reduces the rate of mitotic onset [17]. Akt phosphorylation stabilizes the interaction between Tuberin and Cyclin B1 and as such, permits increased cellular size during energy rich conditions [18].

The ERK signaling pathway is activated downstream of a variety of extracellular stimuli and signals including growth factors, cytokines, G-protein coupled and tyrosine-kinase receptors, integrins and adhesion molecules [1]. These function to relay signals that permit cell growth, proliferation, differentiation, survival, and motility. It has been demonstrated that ERK1 and ERK2 (ERK1/2) phosphorylation on Tuberin occurs at Serine (S) residues 540 and 664, with S644 modifications known to inhibit the binding between Tuberin and Hamartin, leading to an increase of mTOR activity [10]. ERK1/2 phosphorylation can change the subcellular localization of Tuberin, with a phosphorylated mutant presenting a higher nuclear localization when compared to its non-phosphorylated control, alterations which would also reduce Tuberin effects as an inhibitor of mTOR [10]. Clinical studies show TSC-associated brain lesions have constitutively high levels of ERK1/2 activity, with ERK/MAPK inhibitors being suggested as a method of treatment for these TSC-associated brain lesions [19,20]. Moreover, ERK1/2 is hyperactivated in approximately 30% of all cancers with the most common being liver and breast cancers [21,22] and this hyperactivation can support survival of several cancer cell lines [22]. This clinical data supports the idea that understanding the ERK1/2-Tuberin axis will provide important understanding behind TSC driven tumour formation. This study aims to explore the effects of ERK1/2 phosphorylation of Tuberin on cell cycle dynamics.

## Materials and Methods

### Pan-cancer Phosphoproteomic Analysis

Quantitative phosphoproteomic data were obtained from the PhosCancer platform, which integrates mass spectrometry–based phosphoproteomic datasets from the Clinical Proteomic Tumor Analysis Consortium (CPTAC) and related cancer proteomics studies [23]. Abundance values for Tuberin pS664 and RPS6 pS235/S236 were analyzed using protein-corrected normalization as implemented by the platform, whereby phosphosite intensities are adjusted for corresponding total protein abundance. Analyses were performed across tumour cohorts with available phosphoproteomic and clinical annotation, including lung adenocarcinoma (LUAD), lung squamous cell carcinoma (LSCC), glioblastoma (GBM), head and neck squamous cell carcinoma (HNSCC), pancreatic ductal adenocarcinoma (PDAC), clear cell renal cell carcinoma (CCRCC), breast invasive carcinoma (BRCA), and uterine corpus endometrial carcinoma (UCEC).

Differential phosphorylation between tumour and matched normal tissues was assessed within each cohort using the statistical framework implemented in the PhosCancer platform. Overall survival (OS) analyses were conducted using phosphosite abundance values stratified into high and low groups by either cohort-specific median or optimal cutpoint. Survival curves were generated using the Kaplan–Meier method, statistical significance was assessed using the log-rank test, and hazard ratios were estimated using Cox proportional hazards models as implemented by the platform. Median-based stratification was used as the primary approach for cross-cohort comparisons, while cutpoint-based analyses are presented for consistency and exploratory purposes.

To assess whether survival associations observed for Tuberin pS664 reflected canonical mTORC1 signalling output, analogous analyses were performed for protein-corrected phosphorylation of RPS6 at S235 and S236 in lung adenocarcinoma using the same analytical framework. All plots were generated using the PhosCancer platform’s built-in visualization tools.

### Plasmids

pCMV-Tag2-Mock and full-length human pCMV-Tag2-*TSC2* mammalian expression vectors (Flag-tagged) were supplied by J. DeClue. Human full-length CycB1-WT – 5xA and 5xE in a pCMX backbone (GFP tagged) were generous gifts from A. Hagting [24]. Tuberin phosphomimic and unphosphorylated state mutants were constructed using site-directed mutagenesis by the single amino acid change at the S540 and S664 positions.

### Antibodies

The following antibodies were utilized during Western Blotting and indirect immunofluorescent analysis: Mouse α-CycB1 (monoclonal; Santa Cruz; Cat# H1809), mouse α-FLAG (Sigma; Cat# F1804), rabbit β3-Tubulin (monoclonal; NEB; Cat#5568), goat α-mouse IgG (Sigma; Cat# A4416), goat α-rabbit IgG (Sigma; Cat#A6154), Alexa 488-conjugated α-rabbit IgG (Invitrogen; Cat# A-11008), Alexa 568-conjugated α-mouse IgG (Invitrogen; Cat# A-11004), rabbit α-phospho-H3(S-10) (ABcam); Cat# ab32107), RPS6-5G10 (NEB cat #2217) and pRPS6 Ser235/236 (NEB cat# 4858).

### Cell Culture

HEK-293 cells (ATCC) were maintained in Dulbecco’s Modified Eagle Medium (DMEM) supplemented with 10% or 0.5% fetal bovine serum (FBS; Gibco) and 1% Penicillin-Streptomycin (Thermo-Fisher). Cells were incubated at 37 °C in 5% CO2.

### Transfection and Immunoblotting

Sub-confluent HEK-293 cells were transfected with branched PEI (Polysciences, Inc.) 18 to 24 hours after transfection, the cells were lysed or washed with 1X PBS and cultured in normal serum medium (10% FBS) for 24 hours prior to the lysis. The lysed samples were resolved by 10% SDS-PAGE. The membrane was blotted with the indicated antibody (1:1000) followed by enhanced chemiluminescence (ECL) (Amersham). Chemiluminescence was quantified on a ChemiDoc MP V3 (BioRad) and densitometry analysis was performed using BioRad Imaging software. Densitometry is performed by obtaining the total densitometry units of each band. Bands looking at total protein levels (lysate blots) are normalized to the β-tubulin loading control band in the corresponding lane.

### Co-immunoprecipitation

Sub-confluent HEK-293 cells were transfected with branched PEI (Polysciences, Inc.) 18 to 24 hours after transfection, the cells were lysed or washed with 1X PBS and cultured in normal serum medium (10% FBS) for 24 hours prior to the lysis. The lysed samples were then incubated with 1:100 concentrations of Cyclin B1 antibody and overnight at 4°C. The following day, 5µl of Protein G beads were added to the samples and incubated at 4°C for 2 hours. Following this, the samples were centrifuged at 2000 rpm for 2 minutes and supernatant was removed. Fresh lysate buffer was added to each pellet to wash the beads and then centrifuged at 1000 rpm for 2 minutes. The washes were repeated 3 times. The resulting samples were resolved on 10% SDS-PAGE gels as described above.

### Immunofluorescent Microscopy

Cells were seeded onto glass coverslips and transfected as described above. 18 hours following transfection, cells were fixed with 4% formaldehyde in phosphate-buffered saline for 20 minutes and permeabilized with 0.02% Triton X-100 for 5 min. Primary antibodies were used at 1:500, and secondary antibodies at 1:1300. Hoechst stain (Sigma) was added to the permeabilizing solution to a final concentration of 0.5 μg/ml. Slides were imaged by fluorescence microscopy using Zeiss Blue Confocal microscopy with 20x lens and analyzed by the accompanying software.

### Proliferation Assay

HEK-293 cells were seeded onto 6 cm plates and transfected as described above. 16 hours following transfection, the media was removed, and the cells were washed in 1mL of 1X PBS. Fresh media was added to each plate and the cells were incubated for 24, 48 or 72 hours. At each time point, the cells were collected and counted using Trypan Blue to count alive cells. The remaining cells were subjected to cell lysis as described above to prepare for Western blotting.

### Cell Cycle Arrest

Cells were treated with 2 mM of thymidine for 18 hours and then released for 8 hours in full serum media. Then the cells were treated with 2 mM of thymidine for 14 hours. Cells were then collected every two hours for twelve hours after release into full serum media. Cells were then fixed with 70% ethanol and stored at −20°C. Finally, cells were washed and stained with DAPI nuclear stain and subjected to flow cytometry to analyze DNA content of the cells.

### Flow Cytometry Analysis

HEK293T cells were seeded onto 6 cm plates and transfected at sub-confluent numbers with the desired Tuberin mutant. After 24 hours, the cells were collected and washed in 1X PBS. For Rapamycin experiments, the cells were incubated with 100 nM of rapamycin (Selleckchem – S1039) for 4 hrs before collection. The cells were then fixed in 70% ethanol and stored in −20°C. The cells were washed in 1X PBS and then resuspended in PBS-EDTA buffer. DAPI was used for DNA staining. The cells were analyzed using the BD LSRFortessa™ X20 with BD Diva software or FlowJo v10.10.0 software and gated for DNA content.

### Statistical Analysis

Statistical analysis and data visualization were performed using GraphPad Prism (version 10.4.2). Specific statistical tests, number of replicates, and measures of variance are reported in the corresponding figure legends. Data were assessed for significance using appropriate parametric or non-parametric tests, as applicable to each dataset. A *p*-value of <0.05 was considered statistically significant. Error bars represent mean ± SEM, unless otherwise indicated.

## Results

### Pan-cancer analysis reveals heterogeneous ERK-dependent phosphorylation of Tuberin at S664 with context-specific prognostic relevance

To examine the prevalence and clinical relevance of ERK-dependent phosphorylation of Tuberin at S664 in human cancers, we first analyzed pan-cancer phosphoproteomic datasets using the PhosCancer platform. A schematic overview of ERK–Tuberin signalling is shown in Fig 1A. Quantitative analysis of tumour and matched normal tissues revealed that pS664 abundance is not uniformly increased across tumour types but instead displays marked intertumoral heterogeneity (Fig 1B). In several cohorts, bulk tumour pS664 levels were comparable to or lower than those observed in normal tissues, indicating that ERK-mediated phosphorylation of Tuberin is not a universal tumour-wide feature.

**Fig 1.**
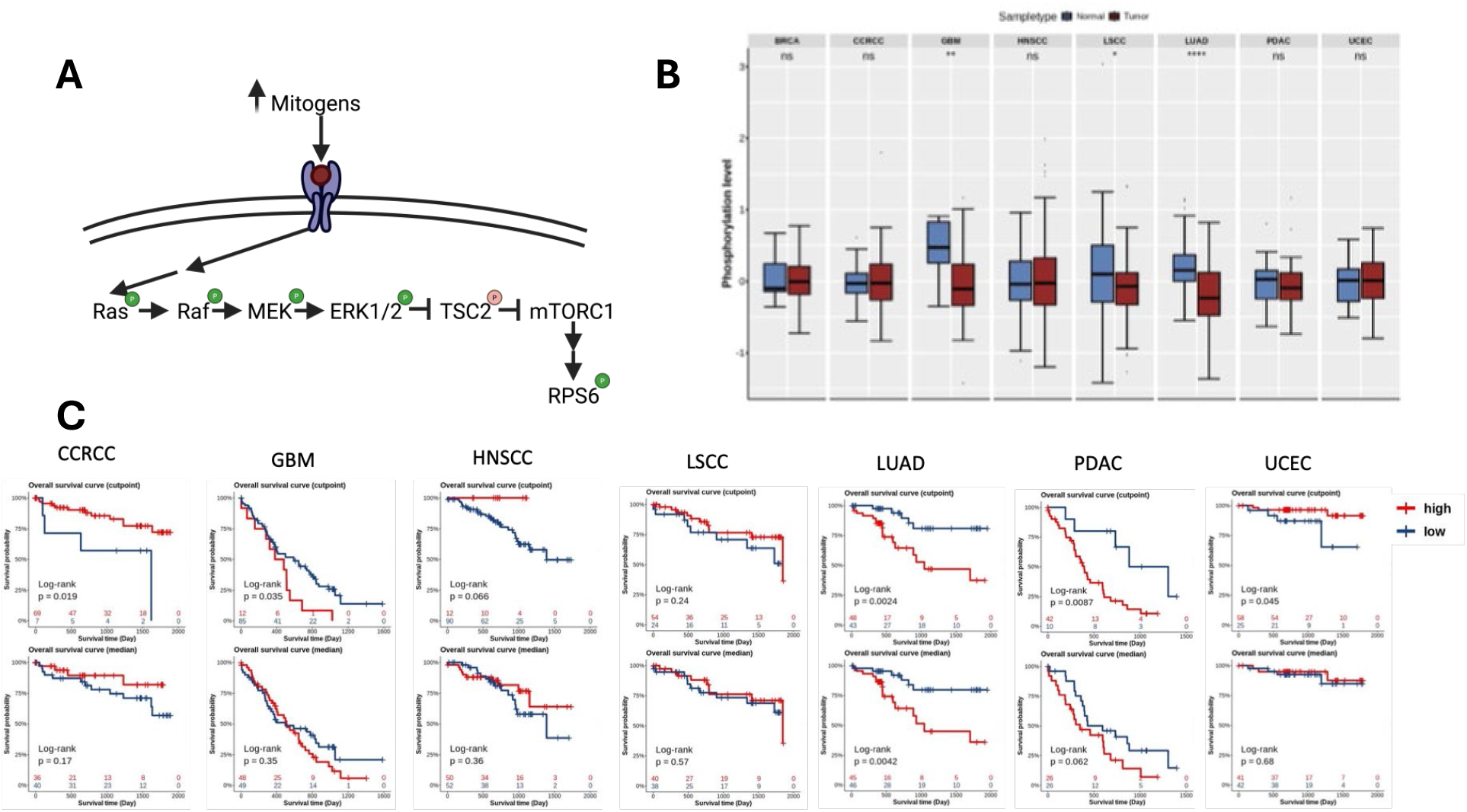
ERK-dependent phosphorylation of Tuberin at S664 is heterogeneous across cancers but associated with adverse outcomes in specific tumour contexts. (A) Schematic of mitogen-activated MAPK signaling illustrating ERK1/2-mediated phosphorylation of Tuberin (TSC2) at S664 and its linkage to downstream mTOR signaling. (B) Differential phosphorylation of TSC2 at S664 in tumour versus matched normal tissues across multiple cancer types, assessed using quantitative phosphoproteomic data from the PhosCancer platform. Phosphorylation levels are shown as protein-corrected phosphosite abundance, highlighting intertumoral heterogeneity rather than uniform tumour-wide upregulation. (C) Overall survival analyses stratified by Tuberin pS664 levels within tumour cohorts. Median-based stratification is shown for all cancer types to enable unbiased cross-cohort comparison, with optimal cut-point analyses included to illustrate subgroup-specific effects. Elevated pS664 is associated with significantly poorer survival in lung adenocarcinoma, consistent with a context-dependent role for ERK-mediated Tuberin phosphorylation.

Despite this heterogeneity in bulk abundance, stratification of tumour cohorts by pS664 levels revealed context-specific associations with clinical outcome. Elevated pS664 was significantly associated with reduced overall survival in lung adenocarcinoma under both median-based and optimal cut-point stratification, whereas no significant survival association was observed in most other tumour types analyzed (Fig 1C). Together, these findings suggest that phosphorylation of Tuberin at S664 marks a biologically distinct and clinically aggressive tumour subset in specific cancer contexts rather than a pan-cancer alteration.

In LUAD, the survival association observed for Tuberin pS664 was not recapitulated by protein-corrected RPS6 pS235/S236 (S1 Fig), a canonical mTORC1/S6K output marker, suggesting that pS664-associated risk is not simply explained by global mTORC1 output and motivating investigation of non-canonical, cell-cycle–linked functions of Tuberin.

### Tuberin pS664 presents reduced affinity for Cyclin B1 when compared to the unphosphorylated form

Tuberin phosphorylation status determines the interaction of this protein with its binding partners [2]. To verify if the Tuberin-Cyclin B1 complex formation is altered by ERK Tuberin phosphorylation, HEK293T cells were transfected with a Flag-tagged Tuberin vector harboring unphosphorylated state Alanine (A) or phosphomimic Aspartic Acid (D) mutations at the ERK phosphorylation sites. These mutations had been previously described in the literature as conferring a phosphomimic and unphosphorylated state [10]. The cells were co-transfected with a mutant form of Cyclin B1, named Cyclin B1 5xA which has been previously described [17]. Cyclin B1 5xA has five Alanine substitutions in the CRS region of Cyclin B1 forcing cytoplasmic localization. Co-immunoprecipitation of Tuberin using Cyclin B1 as a bait shows significantly increase of Tuberin bound to Cyclin B1 when the phosphorylation sites are mutated to Alanine (A) [17]. Our result shows very little change in binding affinity from Tuberin-WT to the mutants expressing unphosphorylated state forms of Tuberin (Fig 2A). Moreover, as is consistent with previous literature [25], the Serine-664 phosphorylation seems to be able to confer the phenotype by itself, regardless of if Serine-540 is substituted to Alanine or Aspartic Acid (Fig 2B). These data support the hypothesis that ERK1/2 phosphorylation of Tuberin alters its ability to bind to Cyclin B1.

**Fig 2.**
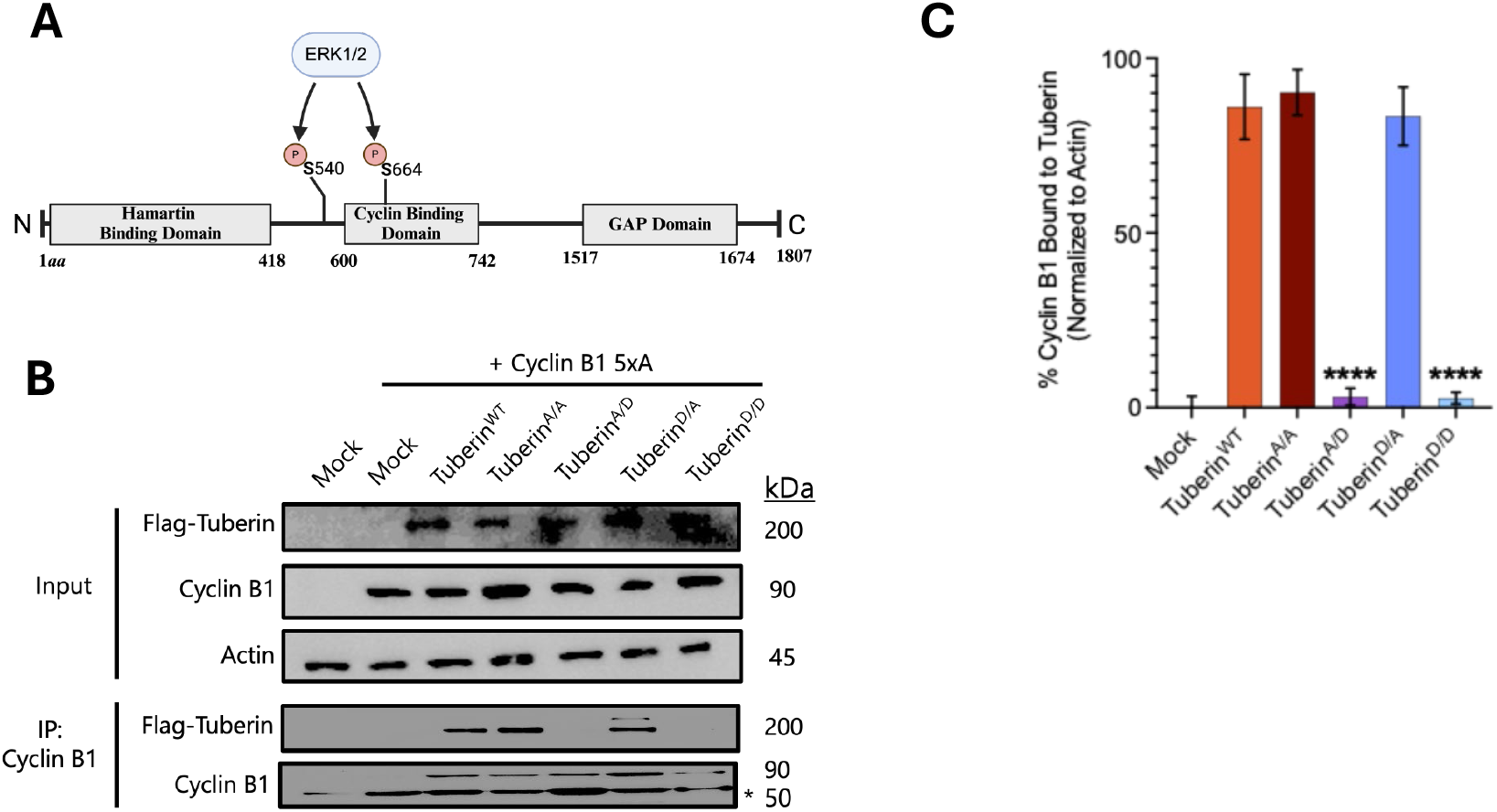
Tuberin p-S664 shows a significant decrease in ability to bind Cyclin B1. (A) Simplified Tuberin primary structure with critical domains reflected. Erk phosphorylation sites on Tuberin, S540 and S664, were targeted using SDM to create phospho-mutant combinations. (B) HEK293T cells were co-transfected with cytoplasmic Cyclin B1 and either empty Flag vector (mock), Flag-Tuberin^WT^ or a phospho-mutant Flag-Tuberin. Lysates were collected after 24 hours and subjected to immunoprecipitation with mouse monoclonal antibody against Cyclin B1. Lysate inputs and immunoprecipitated were subjected to SDS-PAGE and Western blotting. (C) Quantification of the chemiluminescent images measuring densitometry units *n* = 3 biological replicates. One-way ANOVA \*\*\*\**p* < 0.0001.

### Cells overexpressing Tuberin p-S664 show increased proliferation rate

To determine the functional effects of ERK phosphorylation, a proliferation assay was performed. Equal number of sub-confluent HEK293T cells were transfected with mock, Tuberin-WT or Tuberin mutants. A time course proliferation assay was performed where transfected cells were collected 24, 48 or 72 hours post transfection and subjected to trypan blue exclusion. Trypan blue exclusion was used to count live cells at each time point. Tuberin ERK mutant vectors with phosphomimic substitutions at the residue S664 (Tuberin^A/D^ and Tuberin^D/D^) had significantly more cells than the mock control after 72 hours. Whereas the ERK mutants with unphosphorylated state substitutions at S664 (Tuberin^A/A^ and Tuberin^D/D^) showed similar cell numbers to Tuberin-WT and significantly fewer cells compared to mock after 72 hours of growth (Fig 3A). Transfection confirmation was done through western blot (Fig 3B).

**Fig 3.**
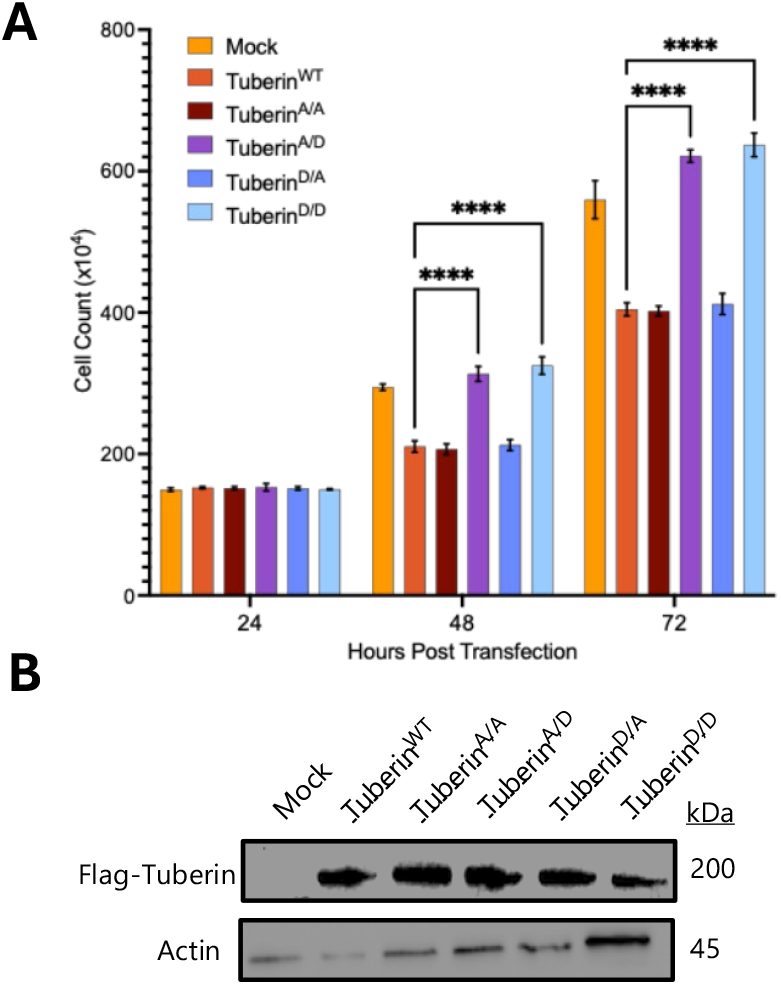
Tuberin p-664 shows increased proliferation. (A) Trypan blue exclusion was used to count live HEK293T cells at 24 hours, 48 hours and 72 hours following transfection with either mock, Flag-Tuberin^WT^ or a phospho-mutant Flag-Tuberin (B) SDS-PAGE and Western blot analysis confirms overexpression of transfected constructs from cell lysates collected at 72 hours; *n* = 3 biological replicates; Two-way ANOVA \*\*\*\**p* < 0.0001.

### Cell cycle profile is altered by Tuberin p-S664 when compared with Tuberin WT

It has been demonstrated that the Tuberin/Cyclin B1 complex delays mitotic onset [17]. To determine the effects of abrogated binding between Tuberin and Cyclin B1 by ERK phosphorylation, flow cytometry was used to assess the cell cycle profile of the Tuberin mutants. HEK293T cells were transfected with a mutant form of Tuberin and subjected to a double thymidine block 24 hours post-transfection. Phosphomimic mutants (Tuberin^A/D^ and Tuberin^D/D^) entered G2 phase faster than the unphosphorylated S-664 state mutants (Tuberin^A/A^ and Tuberin^D/A^), with a significant increase of G2/M cells around 2-4 hours compared to Tuberin^A/A^ and Tuberin^D/A^. Moreover, the Tuberin^A/D^ and Tuberin^D/D^ mutants were able to leave the G2/M phase significantly faster than the Tuberin^A/A^ and Tuberin^D/A^ mutants, with a significant decrease of cells in G2/M phase seen at 10-12 hours for the phosphomimic group (Fig 4).

**Fig 4.**
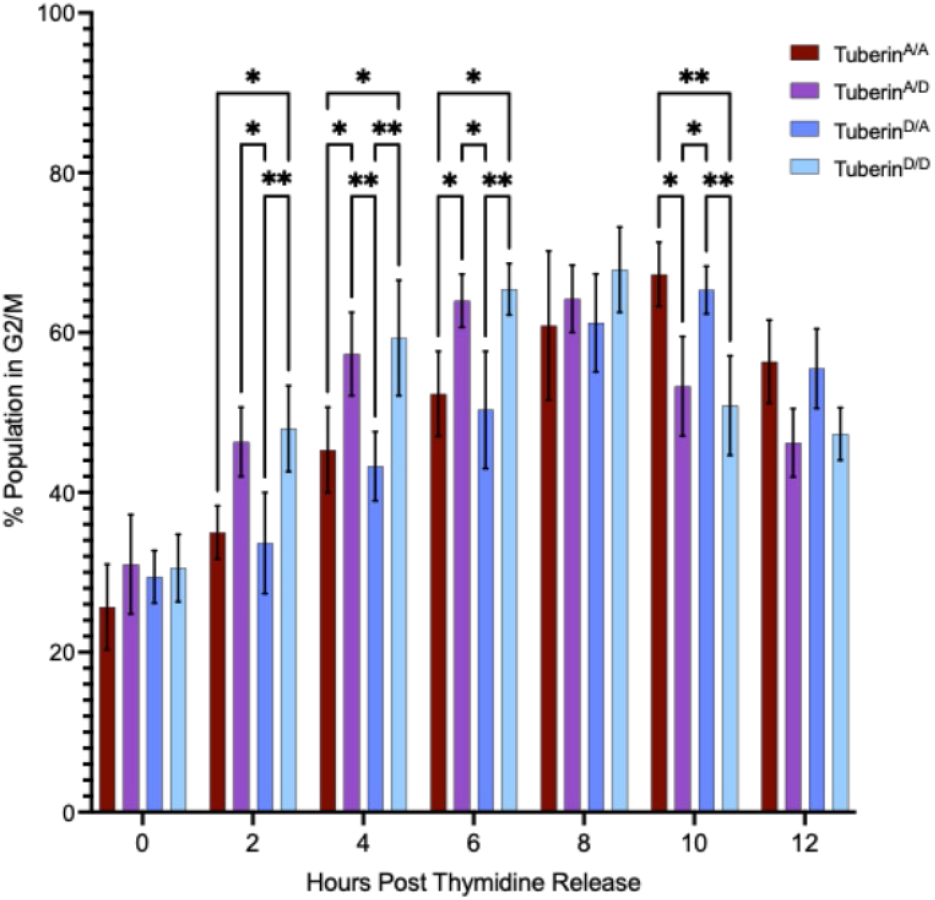
Tuberin Phosphorylated at the ERK sites shows earlier entry into G2/M and exit. HEK293T cells were transfected with mutant form of Flag-Tuberin, and 24 hours post transfection were subjected to a double thymidine block. Following recovery, synchronized cells were collected every 2 hours up to 12 hours, fixed, and stained with DAPI to identify cell cycle profiles using flow cytometry. The percentage of cells from the gated population showed DNA content consistent with cells in the G2/M phase. Tuberin phospho-mutants were statistically compared to each other to determine differences in timing between the different mutants and the time spent in the G2/M phase of the cell cycle, *n* = 3 biological replicates; Two-way ANOVA; * *p* < 0.05, \*\**p* < 0.01.

### ERK Phosphorylation of Tuberin increases mitotic index

Phosphorylation of Histone H3 (pHH3) is widely used to assess mitotic index in cells [26]. To determine the mitotic index of the abrogated binding between Tuberin and Cyclin B1, cells were stained with pHH3 and counted to determine the number of pHH3 positive cells. HEK293T cells were transfected with Tuberin or Tuberin mutants, stained with pHH3 antibody, imaged, and counted for positive cells containing nuclei and green punctate signal for pHH3 positivity. S-664 phosphomimic mutants (Tuberin^A/D^ and Tuberin^D/D^) showed significantly higher pHH3 staining compared to their unphosphorylated state counterparts (Tuberin^A/A^ and Tuberin^D/A^) (Fig 5). Interestingly, the double aspartic acid mutant (Tuberin^D/D^) with phosphorylation at both S540 and S664 residues showed the highest pHH3 staining of the group.

**Fig 5.**
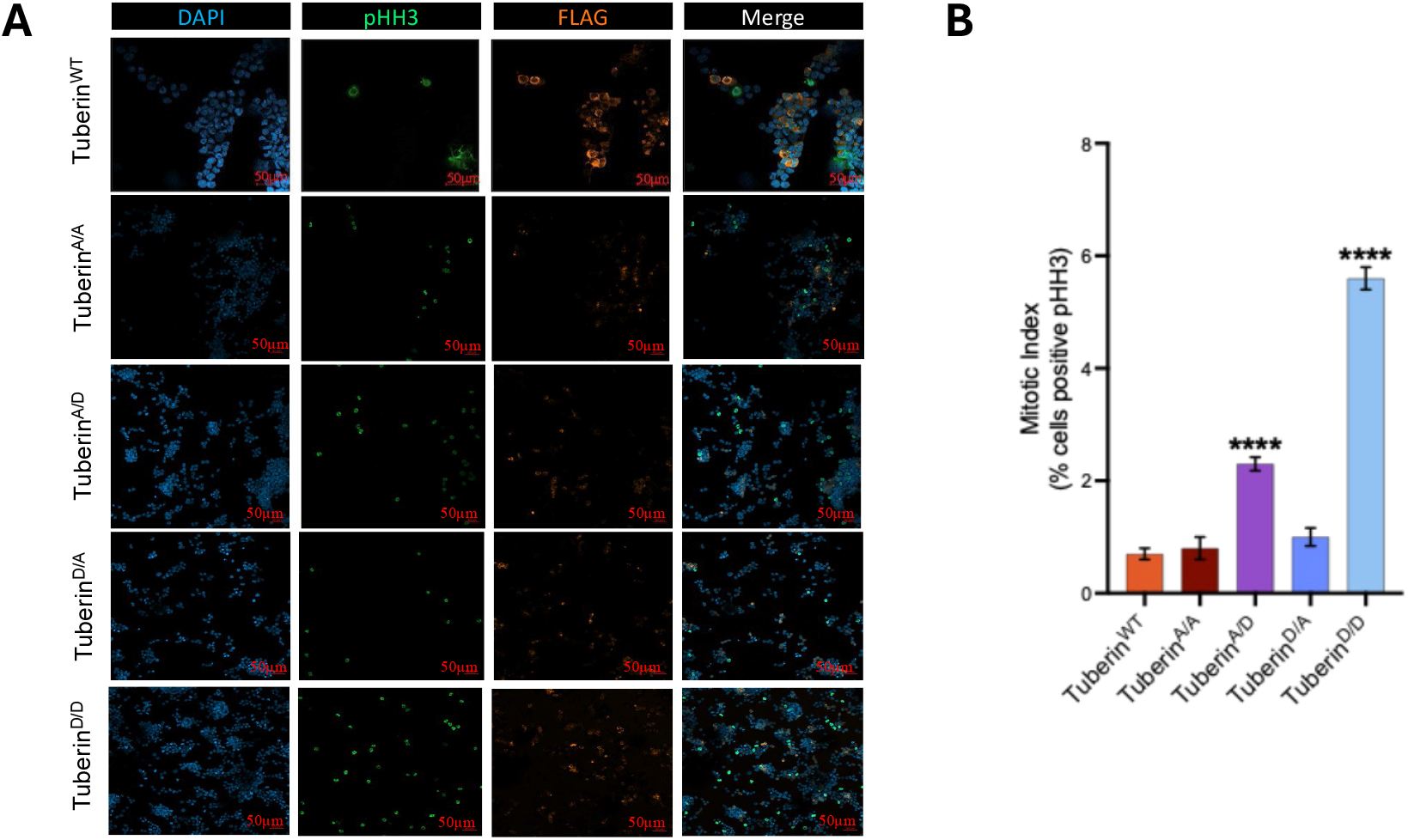
Tuberin phosphorylation at S664 is sufficient to significantly increase mitotic index. HEK293T cells were transfected with Flag-Tuberin^WT^ or phospho-mutant Flag-Tuberin constructs. After 24 hours, coverslips were collected and subjected to immunofluorescent staining. Probing for phospho-histone H3 (pHH3) and Flag was done to score mitotic and Flag-Tuberin expressing cells, respectively. DAPI staining delineates cellular nuclei. (B) Mitotic index scores were calculated based on number of cells that stained positive for pHH3 and Flag; *n =* 3 biological replicates; One-way ANOVA \*\*\*\**p* < 0.0001.

### ERK Phosphorylated Tuberin shows weak co-localization with Cyclin B1

To determine if Tuberin and Cyclin B1 are still localized to the same subcellular compartment, despite having reduced binding potential, cells were co-transfected with Tuberin or Tuberin mutants and Cyclin B1 5xE. This mutant form of Cyclin B1 has glutamate mutations in its CRS region to promote nuclear localization. Cells were grown on coverslips that were imaged on a confocal microscope and images were collected of the cells. The Tuberin^A/D^ and Tuberin^D/D^ mutants show 33% and 16% percent co-localization respectively between Tuberin and Cyclin B1 5xE, and exhibit a punctate staining of Tuberin, typically seen in Tuberin mutants with abrogated binding (Fig 6A and 6B – full panel S2 Fig). These data support that Tuberin^D/D^ has a weak co-localization with Cyclin B1, supporting the abrogated binding seen through co-immunoprecipitation (Fig 2B).

**Fig 6.**
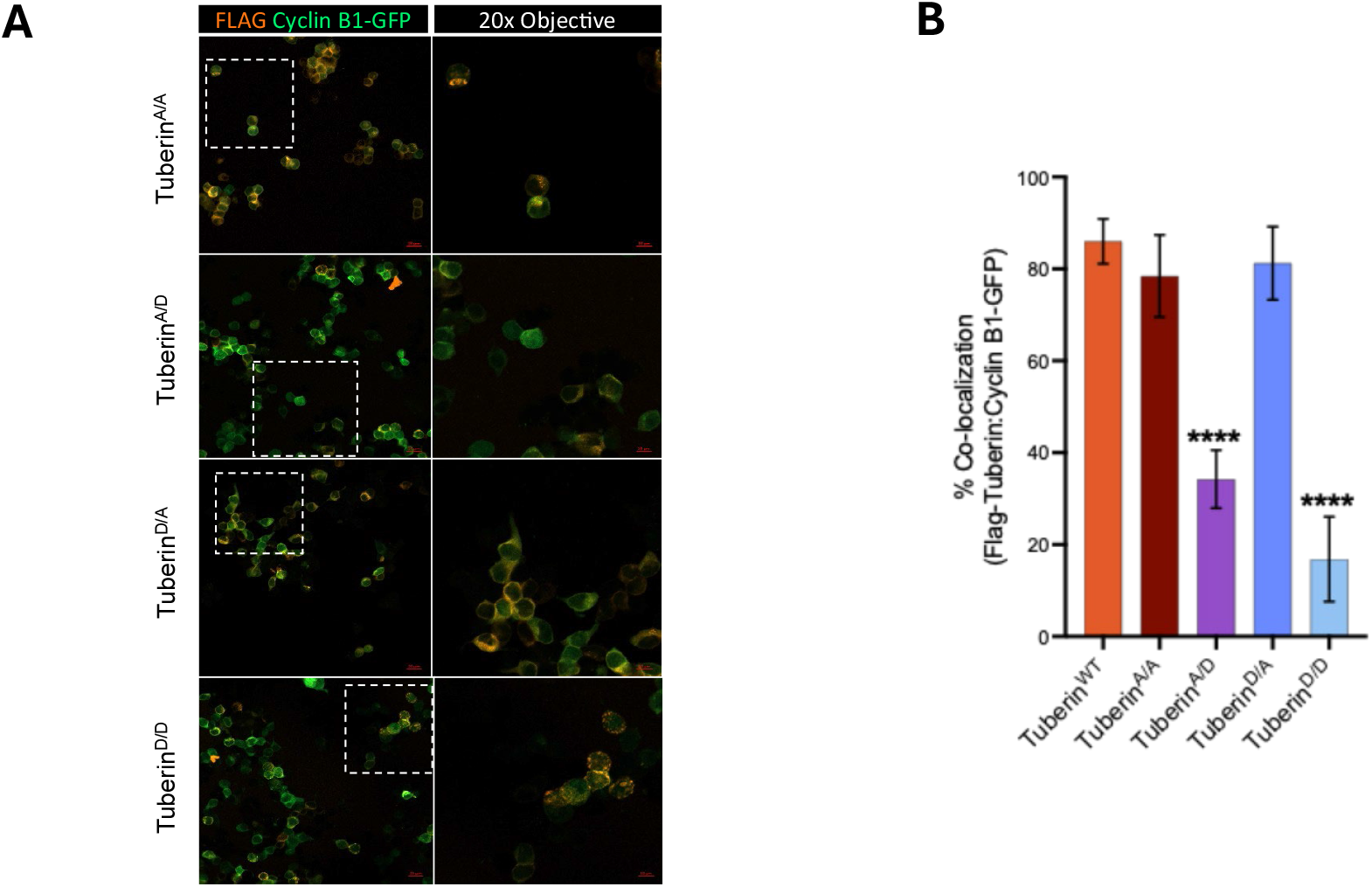
Tuberin Phosphorylated at ERK sites shows weak colocalization with Cyclin B1. (A) HEK293T cells were co-transfected with Tuberin-WT or Tuberin mutant vectors and nuclear Cyclin B1-GFP 5xE. The cells were subjected to immunofluorescent staining for Flag. Regions of interest are set at 20x optical zoom from the confocal field and is outlined in a dotted contour. (B) Quantification of the percent of co-localization, where cells showing significant Flag-Tuberin:Cyclin B1-GFP coalesce were counted and divided by the total number of cells expressing Flag-Tuberin and Cyclin B1-GFP 5xE; *n =* 3 biological replicates. Extended panel in S2 Fig. One-way ANOVA \*\*\*\**p* < 0.0001.

### ERK phosphorylated Tuberin effects in the cell cycle are mTORC1 independent

To test whether cell cycle regulation by Tuberin is independent of mTORC1 we treated the cells with rapamycin, a classic mTORC1 inhibitor, and checked for alteration in the cell cycle profile by flow cytometry (Fig 7). As observed, there isn’t statistically significant difference between the cell cycle profile of cells expressing Tuberin^A/A^ or Tuberin^D/D^ in the presence or absence of rapamycin. These results confirm the Tuberin G2/M arrest isn’t mTORC1 dependent.

**Fig 7.**
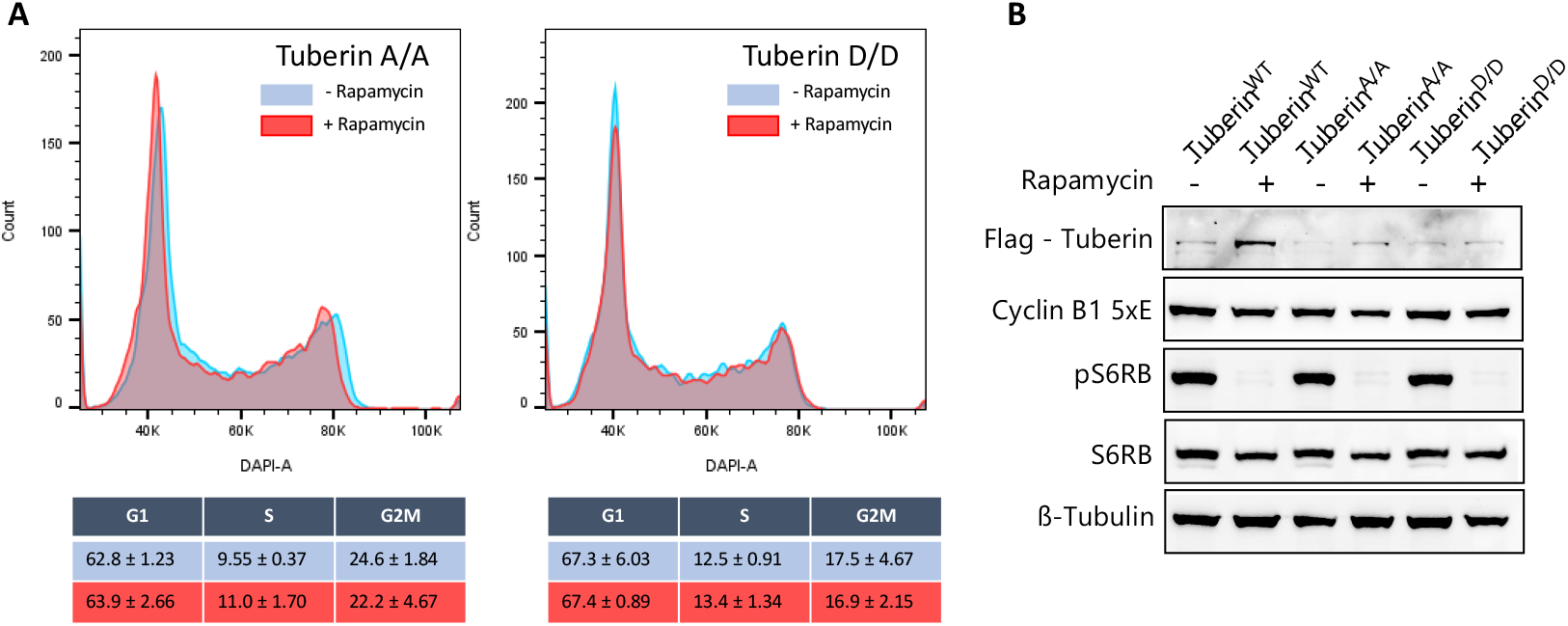
Tuberin G2M arrest is independent of mTORC1 signalling. (A) HEK293T cells were transfected with Tuberin-WT or Tuberin mutant vectors and Cyclin B1-GFP 5xE. After 24 hrs the cells were treated (red) or not (blue) with Rapamycin 100 nM for 4 hrs before collection. Cells were analyzed for the cell cycle profile using DAPI. (n=3) p>0.05. Student’s t-test was performed for each cell cycle phase in the absence or presence of rapamycin, p>0.05 was obtained confirming no statistically significant difference. (B) Western-blot showing non-phospho RPS6 protein in cells treated with rapamycin (+) in opposition to the phosphorylation of RPS6 in absence of rapamycin (-).

## Discussion

Constitutive ERK activation is found as a driving force in a wide variety of cancer types, specifically found in liver and breast cancers [21,22]. The implications of mutations in the ERK pathway are numerous and can be attributed to the many interacting partners found downstream of ERK activation [27]. One of these downstream targets of ERK is Tuberin, a major component of the Tuberous Sclerosis Complex. Our findings refine previous observations of ERK-dependent Tuberin phosphorylation by demonstrating that S664 phosphorylation is not a uniform feature of human cancers, but instead occurs in a heterogeneous, context-dependent manner. This heterogeneity suggests that the biological consequences of S664 phosphorylation may be restricted to specific tumour states, motivating focused mechanistic studies. The canonical function of this complex is to regulate the activity of protein synthesis through its control of the mTOR pathway [28]. ERK regulates the activity of the TSC complex through its kinase activity towards Tuberin at the phosphorylation sites at S540 and S664. Phosphorylation has previously been shown to dissociate Tuberin and Hamartin, the other major component of the TSC, allowing for the degradation of Tuberin and activation of mTOR and protein synthesis [10,29,30]. Tuberin is also a regulator of the G2/M checkpoint through an interaction with Cyclin B1, wherein, Tuberin retains Cyclin B1 in the cytoplasm to delay mitotic onset [17]. Much less is known about the regulation of this interaction. Our previous work demonstrated that Akt phosphorylation of Tuberin occurring downstream of metabolic pathways including insulin signaling increased binding between Tuberin and Cyclin B1 [18]. This work building on this knowledge by demonstrating that growth factor mediated ERK1/2 pathway abrogates the binding between Tuberin and Cyclin B1 through the phosphorylation of S664 on Tuberin in mTORC1 independent pathway.

## Conclusion

The actual mechanism surrounding the G2/M transition in the cell cycle is historically understudied when compared to other aspects of the cell cycle [31]. The understanding of how cells can recognize the extra-cellular environment and integrate signals into the decision to progress from G2 phase into mitosis is crucial to understand proper cell division and mechanisms of proliferative diseases. The results found in this study, combined with previous studies from our group and others [16–18] further illuminate how the G2/M transition is regulated by nutrients and growth factors. It can be hypothesized that activation of ERK1/2 and inhibition of Akt signalling [18] present a situation where Cyclin B1 can easily translocate to the nucleus thereby enhancing cell proliferation (Fig 8). This may have important mechanistic relevance for cancer initiation pathways where ERK activation is a common change [1]. While the mechanistic insights presented here are derived from controlled *in vitro* systems, they provide a framework for understanding how ERK-dependent phosphorylation of Tuberin may influence G2/M regulation in defined cellular contexts. Extending these findings to *in vivo* models will be important to determine how such signalling dynamics operate within the complexity of tumour microenvironments.

**Fig 8.**
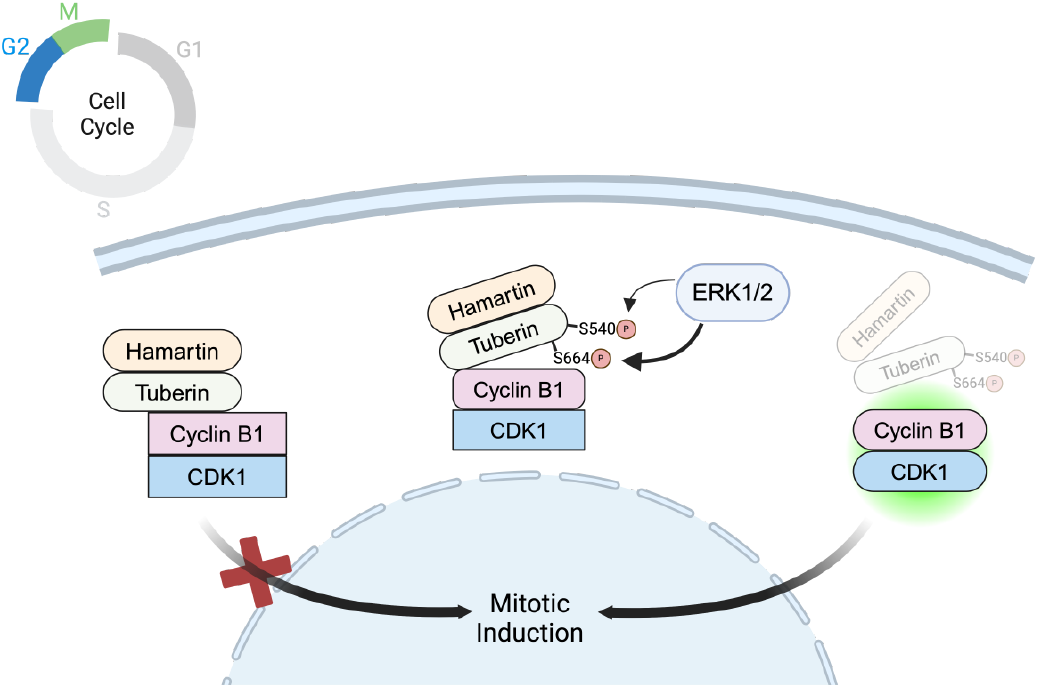
Tuberin binding to Cyclin B1 and G2/M regulation is influenced by the ERK phosphorylation site S664. Unphosphorylated Tuberin holds Cyclin B1/CDK1 complex in the cytoplasm. Upon ERK phosphorylation, post-translational modification in Tuberin allows Cyclin B1/CDK1 translocation to the nucleus.

## Acknowledgement

Our work benefits from the support and input of all members of the Porter Lab, we are grateful for the diversity of contributions. We thank the Imaging Facility at the University of Windsor.

