## Supplementary material for "Effects of ERK1/2 Signaling on Cell Cycle Regulation by the Tuberin-Cyclin B1 Complex": S1 Fig

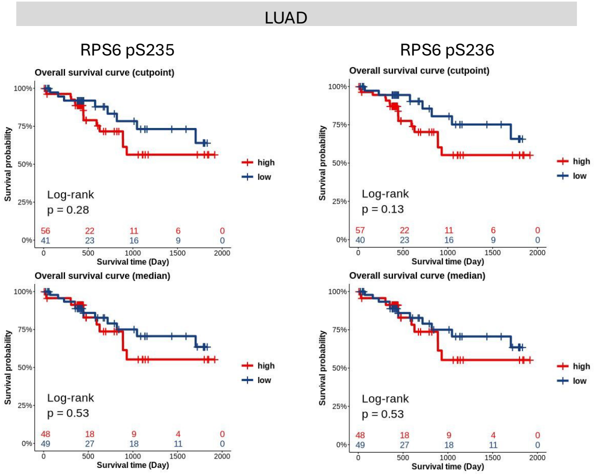


**S1 Fig. Canonical mTORC1 output does not recapitulate the LUAD survival association observed for Tuberin pS664.** Overall survival in lung adenocarcinoma (LUAD) stratified by protein-corrected phosphorylation of RPS6 at S235 and S236. Survival curves are shown using both optimal cut-point and median-based stratification for consistency with the analyses in Fig 1. Under neither stratification approach did RPS6 phosphorylation significantly associated with overall survival, indicating that the prognostic signal linked to Tuberin pS664 in LUAD is not explained by canonical mTORC1/S6K output.
