## Supplementary material for "Effects of ERK1/2 Signaling on Cell Cycle Regulation by the Tuberin-Cyclin B1 Complex": S2 Fig

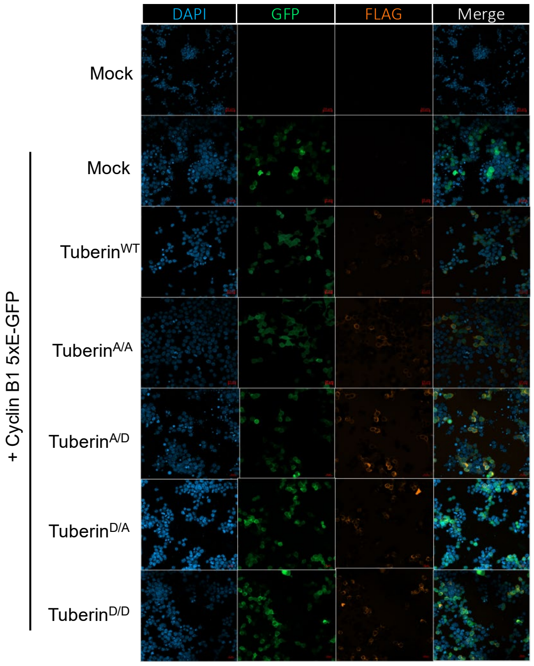


**S2 Fig. Full Panel of Tuberin Phosphorylated at ERK sites shows weak colocalization with Cyclin B1.** HEK293T cells are co-transfected with Tuberin-WT or Tuberin mutant vectors and Cyclin B1-GFP 5xE. After 24 hours, coverslips are collected and subjected to immunofluorescent staining.
